## Supplementary material for "Hsf1 and the molecular chaperone Hsp90 support a “rewiring stress response” leading to an adaptive cell size increase in chronic stress": Figure supplements 1 to 13, Tables 1 and 2, Description of the contents of the files Source Data 1 to 4

**This PDF file includes:**

Figure supplements 1 to 13
Tables 1 and 2
Description of the contents of the files Source Data 1 to 4

**Other Supplementary Materials for this manuscript include the following:**

Source Data 1 to 4

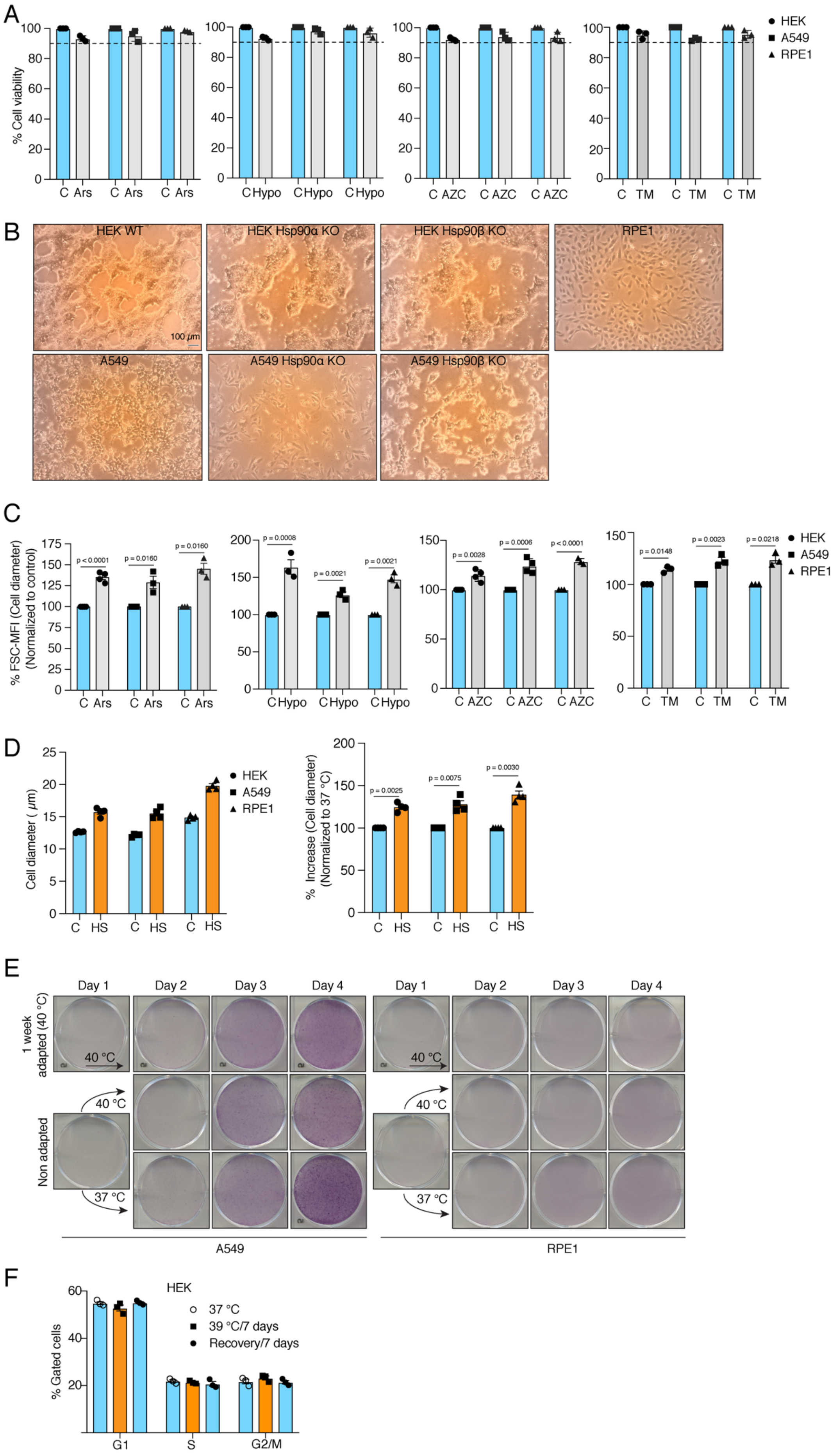
A
% Cell viability
C Ars C Ars C Ars
C Hypo C Hypo C Hypo
C AZC C AZC C AZC
C TM C TM C TM
HEK
A549
RPE1
B
HEK WT
HEK Hsp90α KO
HEK Hsp90β KO
RPE1
100 μm
A549
A549 Hsp90α KO
A549 Hsp90β KO
C
% FSC-MFI (Cell diameter) (Normalized to control)
p < 0.0001
p = 0.0160
p = 0.0160
p = 0.0008
p = 0.0021
p = 0.0021
p = 0.0028
p = 0.0006
p < 0.0001
p = 0.0148
p = 0.0023
p = 0.0218
HEK
A549
RPE1
D
Cell diameter ( μm)
C HS C HS C HS
HEK
A549
RPE1
% Increase (Cell diameter) (Normalized to 37 °C)
p = 0.0025
p = 0.0075
p = 0.0030
E
Day 1
Day 2
Day 3
Day 4
1 week adapted (40 °C)
Non adapted
40 °C
37 °C
A549
RPE1
F
HEK
37 °C
39 °C/7 days
Recovery/7 days
% Gated cells
G1
S
G2/M

**Figure supplement 1. Cells increase their size in response to different types of chronic stress. (A)** Flow cytometric quantification of cell viability after 4 days of treatment with 10 μM sodium arsenite (Ars), 1% hypoxia (Hypo), 5 μM L-azetidine-2-carboxylic acid (AZC), or 250 nM tunicamycin (TM) (n = 3 biologically independent samples). **(B)** Representative phase-contrast micrographs showing the confluency of the indicated cells on the day of harvesting. The size bar in the first macrograph (top left) is 100 μM and applies to all micrographs. **(C)** Flow cytometric quantification of cell size after 4 days of treatment as indicated (n = 4, 3, 4, and 3 biological replicates for Ars, Hypo, AZC, and TM, respectively, for HEK cells; n = 3, 3, 4, and 3 for Ars, Hypo, AZC, and TM, respectively, for A549 cells; n = 3, 3, 3, and 3 for Ars, Hypo AZC, and TM, respectively, for RPE1). **(D)** Bar graphs of the measurements of cell diameter by automated cell counter (on the left) and the percent change in cell diameter (on the right) **(E)** Scanned images of plates with crystal violet-stained cells (representative images of n = 3 independent experiments). **(F)** Flow cytometric analysis of cell cycle after 1 week in chronic HS and post HS recovery (n = 3 biologically independent samples). The data are represented as mean values ± SEM for all bar and line graphs. The statistical significance between the groups was analyzed by two-tailed unpaired Student's t-tests.

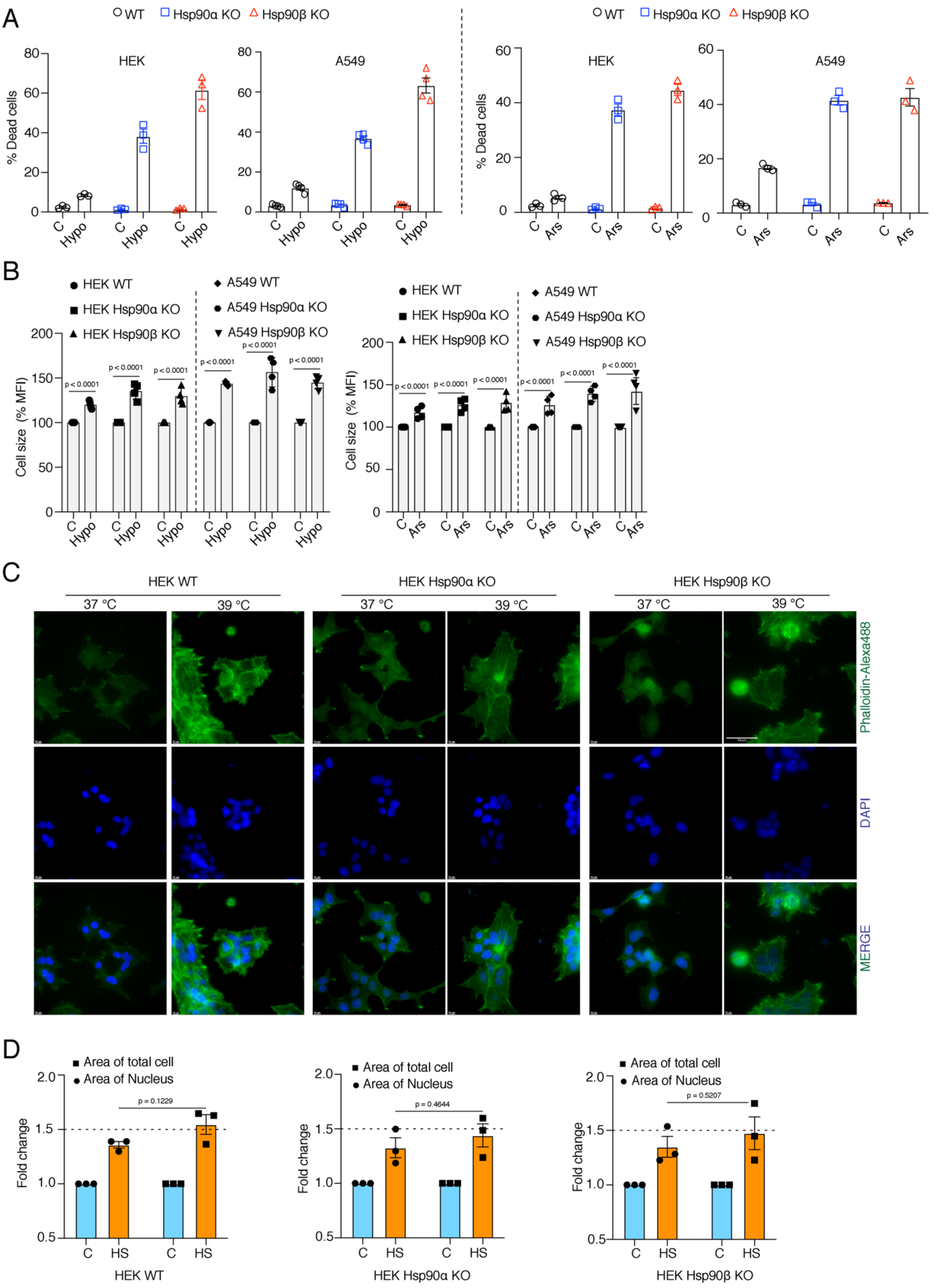
A
WT
Hsp90α KO
Hsp90β KO
HEK
A549
% Dead cells
C
Hypo
Ars
B
HEK WT
HEK Hsp90α KO
HEK Hsp90β KO
A549 WT
A549 Hsp90α KO
A549 Hsp90β KO
Cell size (% MFI)
p < 0.0001
C
HEK WT
HEK Hsp90α KO
HEK Hsp90β KO
37 °C
39 °C
Phalloidin-Alexa488
DAPI
MERGE
D
Area of total cell
Area of Nucleus
Fold change
p = 0.1229
p = 0.4644
p = 0.5207
HS

**Figure supplement 2. Cells are unable to adapt to chronic stress in the absence of one of the Hsp90 isoforms, but still get larger. (A)** Flow cytometric quantification of cell death of A549 WT and Hsp90α/β KO cells after 4 days in 1% hypoxia (Hypo) and 10 μM sodium arsenite (Ars) treatment (n = 3 biologically independent samples). **(B)** Flow cytometric quantification of cell size after 4 days in 1% hypoxia (Hypo) and 10 μM sodium arsenite (Ars) treatment (n = 4 biologically independent samples). **(C)** Fluorescence microscopy images of HEK WT and Hsp90α/β KO cells after 4 days of chronic HS. The cytoskeleton is stained with Phalloidin-Alexa488 (green), and the nucleus is stained with DAPI (blue). The scale bars on each image are 10 μM, and 50 *μ*M on the image at the top right. **(D)** Bar graphs of the area of nuclei and whole cells determined from fluorescent micrographs (representative images are shown in panel C) using ImageJ. We used micrographs from 3 biologically independent experiments. From each experiment we measured 30 randomly chosen cells, and their average values were used as one data point**.** The data of all bar graphs in the figure are represented as mean values ± SEM for all bar graphs. The statistical significance between the groups was analyzed by two-tailed unpaired Student's t-tests.

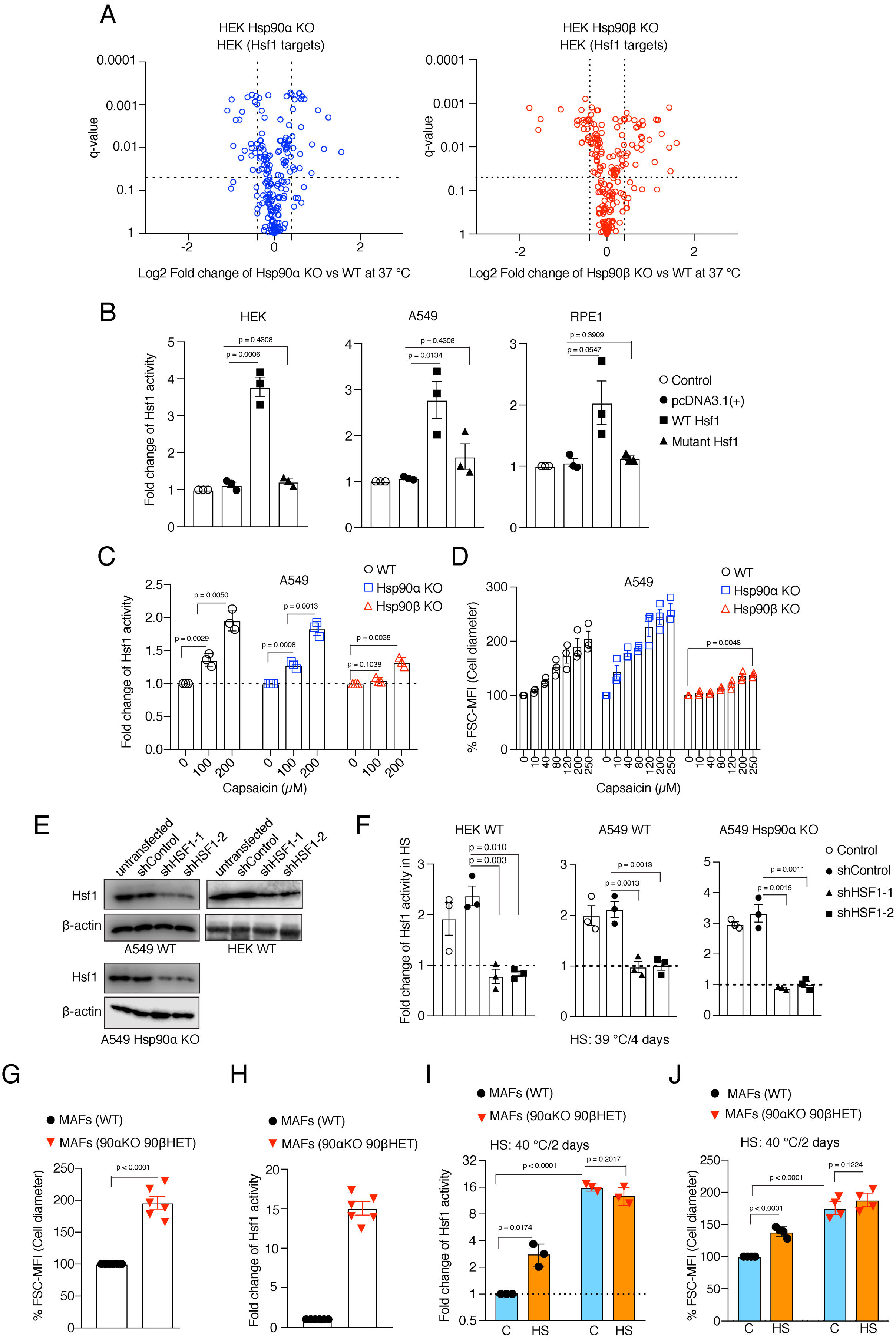
A
HEK Hsp90α KO
HEK (Hsf1 targets)
q-value
Log2 Fold change of Hsp90α KO vs WT at 37 °C
HEK Hsp90β KO
HEK (Hsf1 targets)
Log2 Fold change of Hsp90β KO vs WT at 37 °C
B
HEK
A549
RPE1
Fold change of Hsf1 activity
Control
pcDNA3.1(+)
WT Hsf1
Mutant Hsf1
C
A549
WT
Hsp90α KO
Hsp90β KO
Capsaicin (μM)
D
% FSC-MFI (Cell diameter)
E
untransfected
shControl
shHSF1-1
shHSF1-2
Hsf1
β-actin
A549 WT
HEK WT
A549 Hsp90α KO
F
HEK WT
A549 WT
A549 Hsp90α KO
Fold change of Hsf1 activity in HS
HS: 39 °C/4 days
G
MAFs (WT)
MAFs (90αKO 90βHET)
H
I
HS: 40 °C/2 days
J

**Figure supplement 3. Hsf1 induces cell size in response to stress. (A)** Volcano plots of the normalized fold changes in protein levels of some of the core Hsf1 target genes (list obtained from https://hsf1base.org/) in Hsp90α/β KO cells compared to WT HEK as determined by quantitative label-free proteomic analysis. Molecular chaperones, whose expression is regulated by Hsf1, are excluded from this dataset (n = 3 biologically independent samples). Log2 fold changes of > 0.5 or < -0.5 with a q-value (adjusted p-value) of < 0.05 were considered significant differences for a particular protein. **(B)** Fold change of Hsf1 activity of HEK, A549, and RPE1 cells upon overexpressing WT and mutant Hsf1 in combination with EGFP, as measured with the Hsf1 luciferase reporter. Control is transfected with only Hsf1 reporter plasmid and pEGFP-C1, those are common to all the experimental conditions; (n = 3 biologically independent samples). **(C)** Fold change of Hsf1 activity in A549 WT, Hsp90α KO, and Hsp90β KO cells after 4 days of capsaicin treatment as measured by luciferase reporter assay (n = 3 biologically independent samples). **(D)** Flow cytometric quantification of cell size after 4 days of capsaicin treatment of Hsp90α/β KO and WT A549 cells (n = 3 biologically independent samples). **(E)** Immunoblots of Hsf1 after Hsf1 knockdown in A549 WT, A549 Hsp90α KO, and HEK WT cells. β-actin serves as the loading control (representative of n = 2 independent experiments). **(F)** Fold change of Hsf1 activity in HEK WT, A549 WT, and A549 Hsp90α KO cells in chronic HS after Hsf1 knockdown as measured by luciferase reporter assays. Here the chronic HS for A549 cells is 39 °C to instead of 40 °C to reduce HS-induced damage in Hsf1 knockdown conditions (n = 3 biologically independent samples). **(G)** Flow cytometric quantification of cell size of mouse fibroblast. (90αKO, 90β HET), homozygous *hsp90α* KO, heterozygous *hsp90β* KO cells (n = 6 biologically independent samples). **(H)** Fold change of Hsf1 activity in (90αKO, 90β HET) MAFs compared to WT at 37 °C, as measured by luciferase reporter assays (n = 3 biologically independent samples). **(I)** Fold change of Hsf1 activity in MAFs subjected to chronic HS (orange bars) compared to 37 °C (blue bars), as measured by luciferase reporter assay (n = 3 biologically independent samples). **(J)** Flow cytometric quantification of cell size of MAFs subjected to chronic HS (orange bars) by comparison to 37 °C (blue bars) (n = 4 biologically independent samples). The data are represented as mean values ± SEM for all bar graphs. The statistical significance between the groups was analyzed by two-tailed unpaired Student's t-tests.

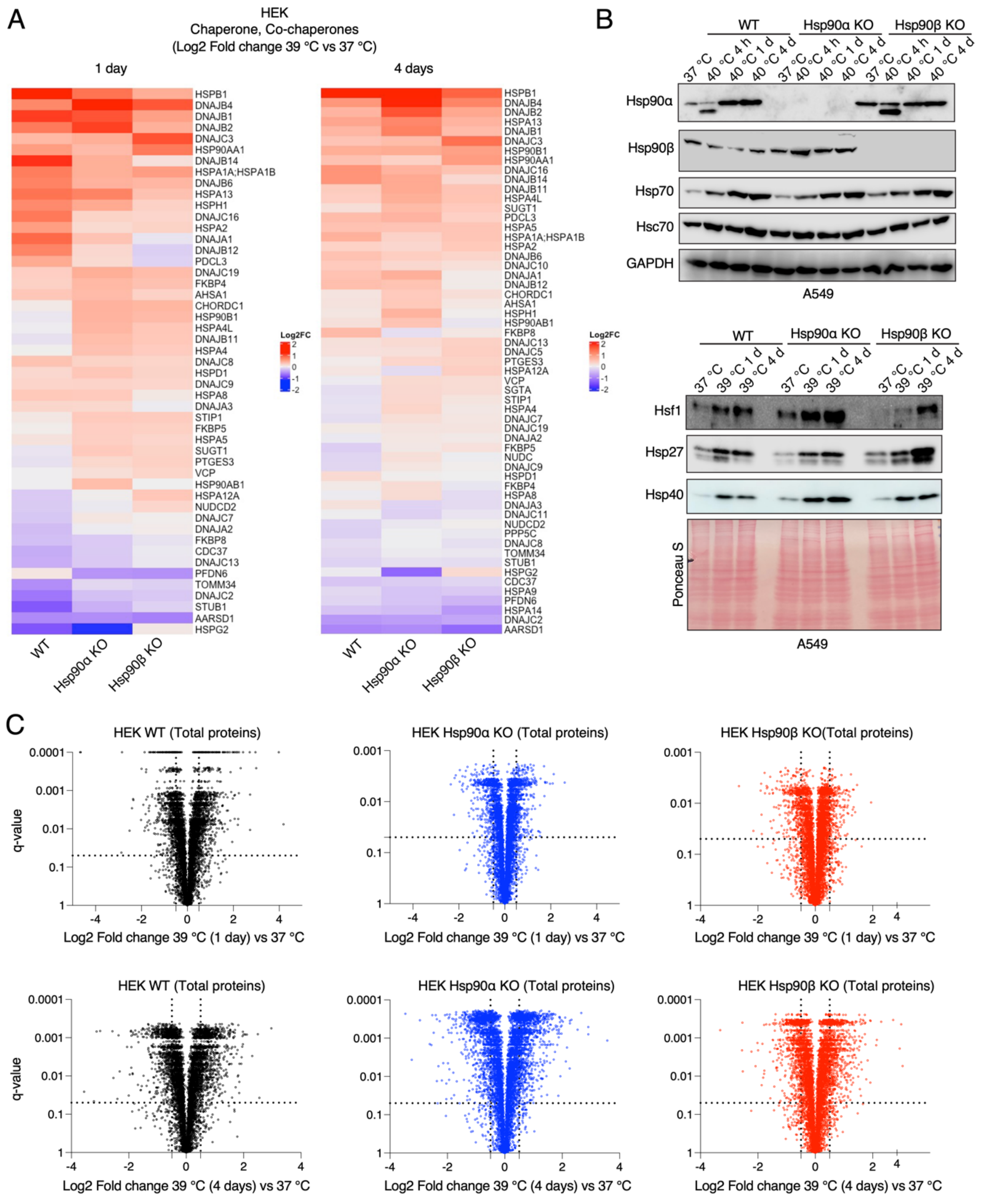


**Figure supplement 4. Hsp90α/β KO cells maintain molecular chaperones, co-chaperones, and total proteins. (A)** Heat maps of the normalized fold changes (log2) of molecular chaperones and co-chaperones of HEK cells subjected to 1 and 4 days of chronic HS as determined by quantitative label-free proteomics (n = 3 biologically independent samples); only proteins whose abundance changed with q-values (adjusted p-values) of <0.05 were considered significant and are shown in the heat map. **(B)** Immunoblots of some molecular chaperones of A549 WT and Hsp90α/β KO cells. GAPDH and the Ponceau S-stained nitrocellulose filter serve as loading controls. **(C)** Volcano plots of the normalized fold changes of total proteins of cells subjected to chronic HS for 1 and 4 days (first and second rows, respectively) determined by quantitative label-free proteomics. Each genotype was compared with its respective 37 °C control (n = 3 biologically independent samples). Log2 fold changes of > 0.5 or < -0.5 with q-values (adjusted p-values) of < 0.05 were considered significant differences for a particular protein.

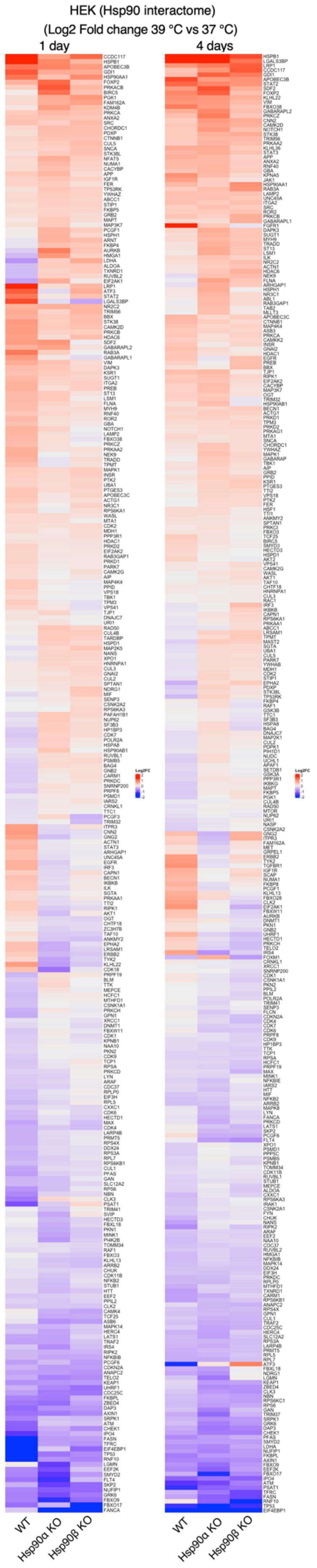
HEK (Hsp90 interactome)
(Log2 Fold change 39 °C vs 37 °C)
1 day
4 days
WT
Hsp90α KO
Hsp90β KO
WT
Hsp90α KO
Hsp90β KO

**Figure supplement 5. Hsp90α/β KO cells maintain Hsp90 interactors in chronic stress.** Heat maps of the normalized fold changes (log2) of Hsp90 interactors of HEK cells subjected to 1 and 4 days of chronic HS as determined by quantitative label-free proteomics (n = 3 biologically independent samples); only proteins whose abundance changed with q-values (adjusted p-values) of <0.05 were considered significant and are shown in the heat map.

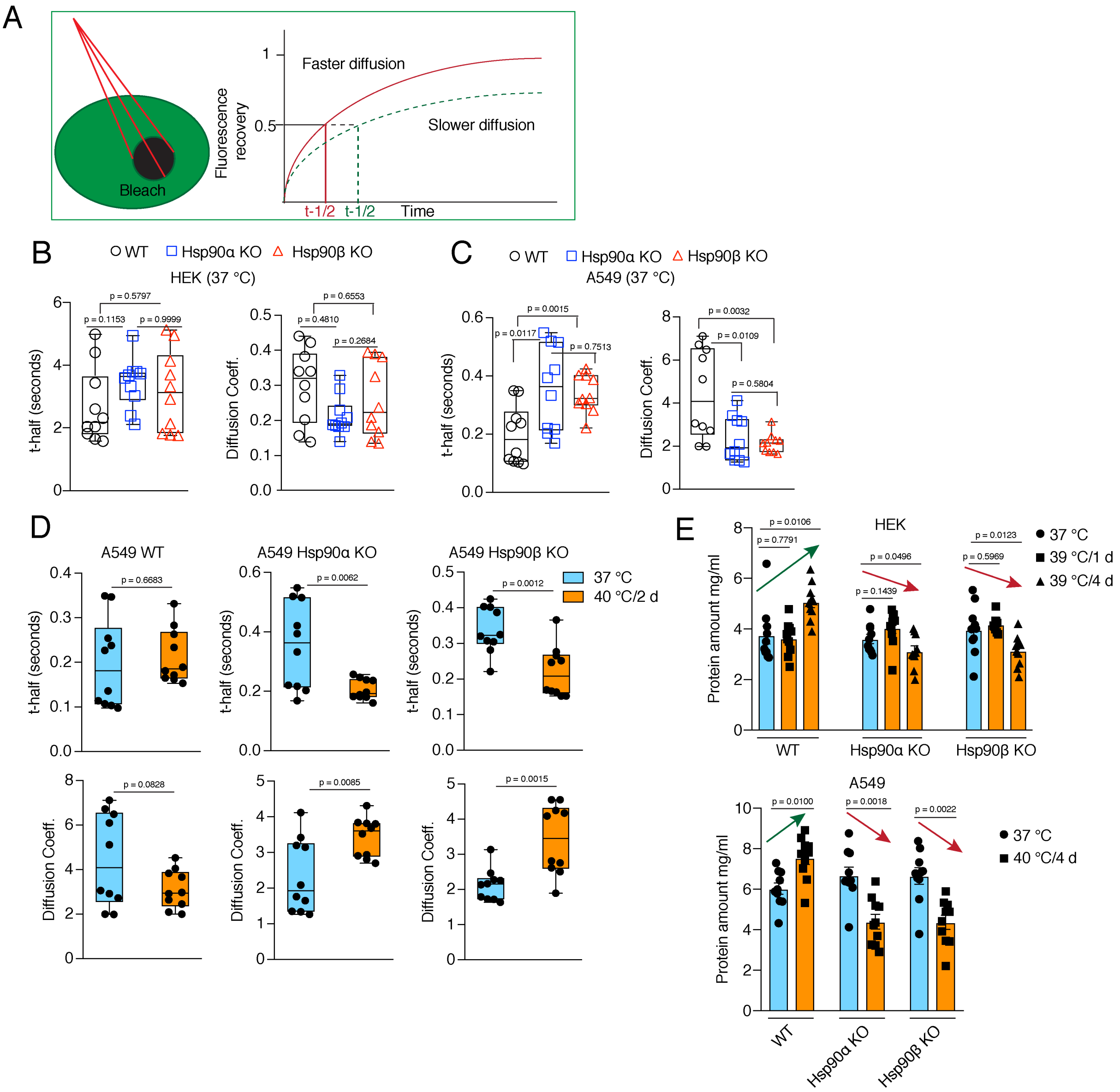


**Figure supplement 6. Wild-type cells maintain cytoplasmic density and total protein ratio during stress-induced cell size increase. (A)** Scheme of the FRAP experiments. **(B to D)** FRAP experiments with live cells expressing EGFP. The respective box plots represent the t-half values of recovery of EGFP fluorescence and the apparent EGFP diffusion coefficients (n= 10 cells from 2 biologically independent experiments). **(E)** Protein amount in cell lysates (5,000 cells lysed in 200 µl lysis buffer) of control cells and cells adapted to chronic HS as measured by Bradford assay.

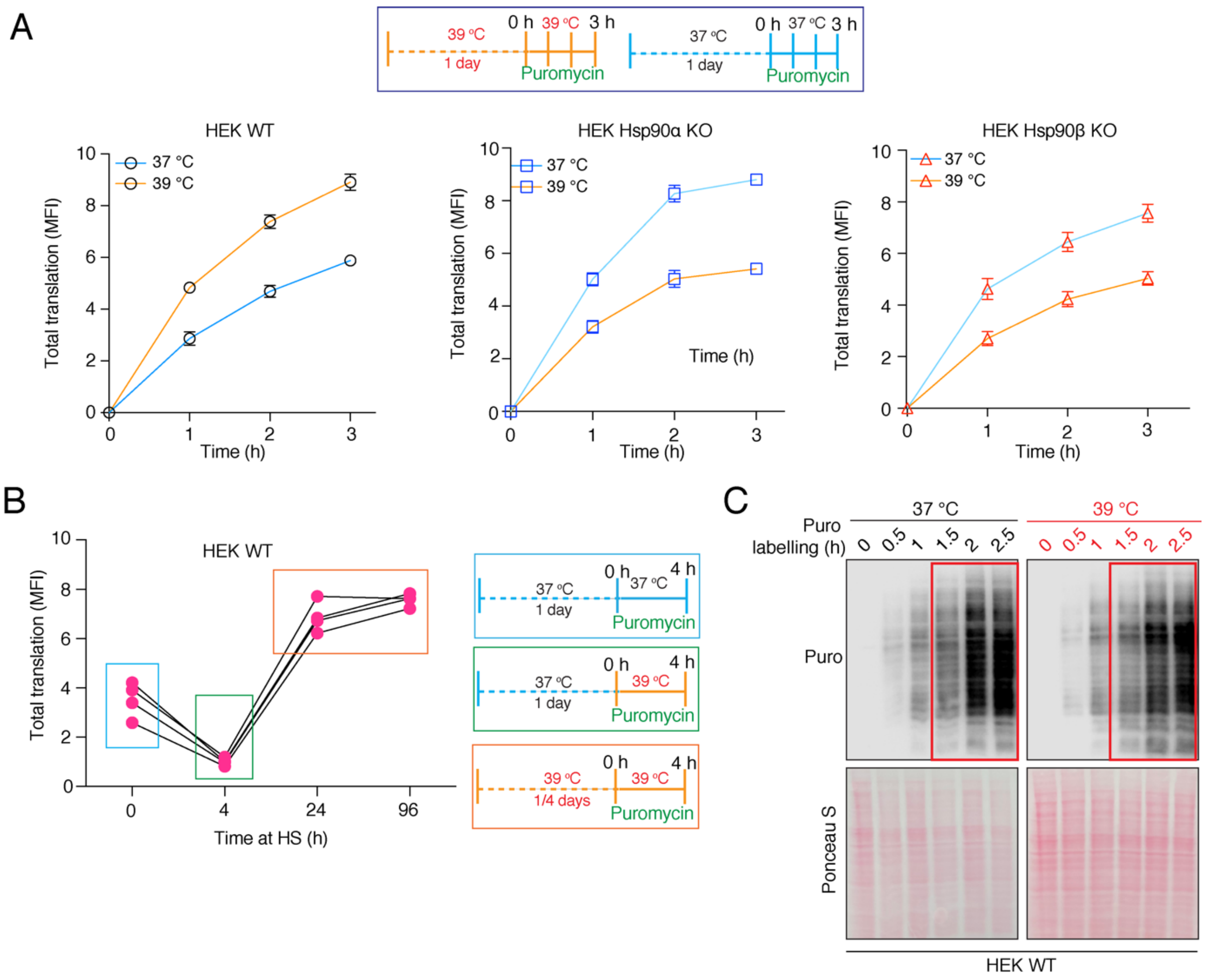


**Figure supplement 7. Hsp90 requirement for cellular translation during adaptation to chronic stress, and early time points of translational adaptation of wild-type cells. (A)** Flow cytometric analysis of global translation of HEK WT and Hsp90α/β KO cells at 37 °C and after 1 day under chronic HS. See scheme of the experiment on the top. Nascent polypeptide chains were labeled with OP-puromycin during cell culture, and the incorporation of puromycin at different time points was analyzed (n = 4 experimental samples). **(B)** Total translation of HEK WT cells during a 4 h time span during different phases of chronic HS**. (C)** Immunoblot analysis of global translation as indicated by incorporation of puromycin into nascent polypeptides during the first 2.5 h of shifting cells to chronic HS conditions, compared to cells remaining at 37 °C. The 0 h time point of puromycin labelling serves as a negative control, and the Ponceau S-stained nitrocellulose filter as loading control (representative images from n = 2 biologically independent experiments)**.**

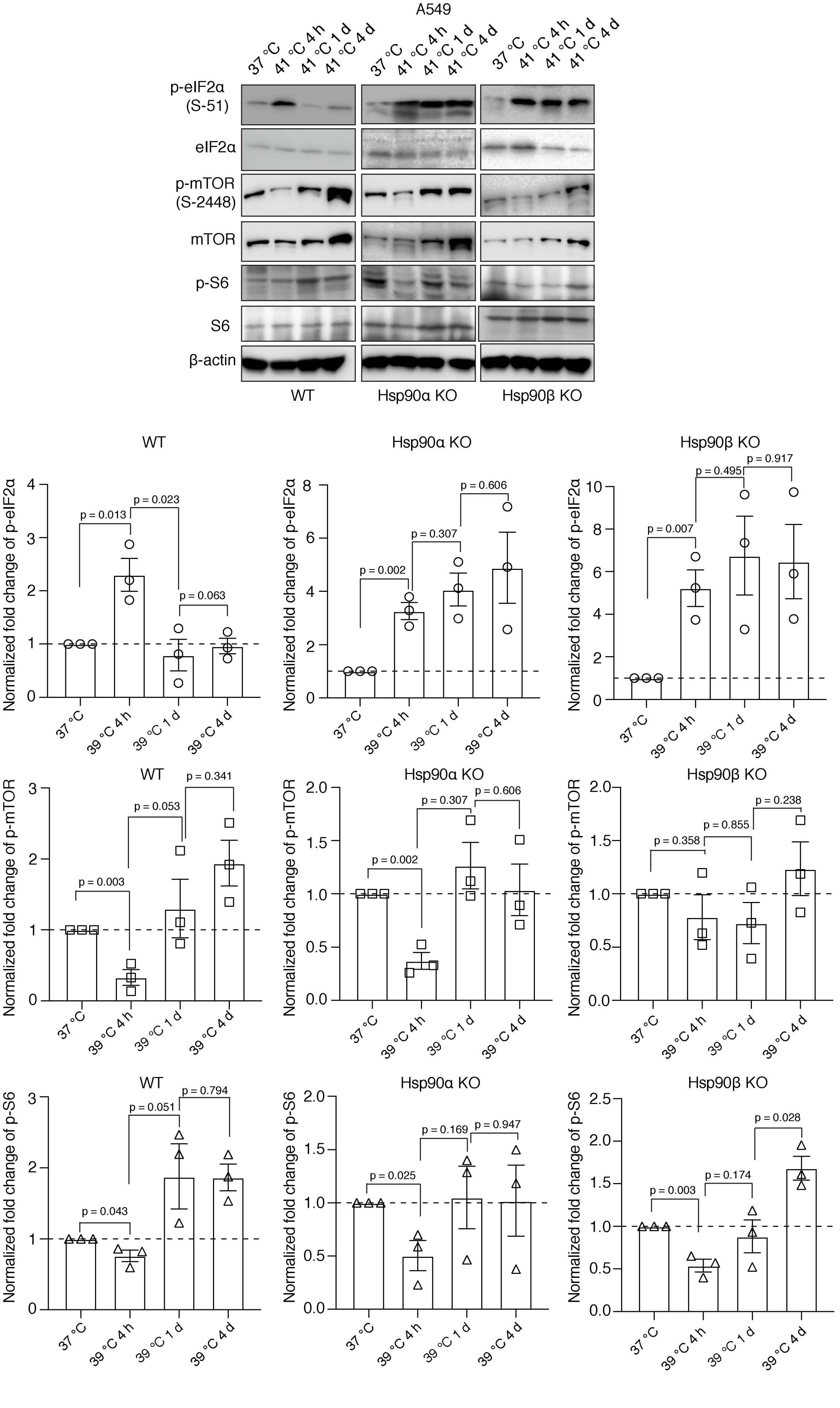
A549
37 °C
41 °C 4 h
41 °C 1 d
41 °C 4 d
p-eIF2α (S-51)
eIF2α
p-mTOR (S-2448)
mTOR
p-S6
S6
β-actin
WT
Hsp90α KO
Hsp90β KO
Normalized fold change of p-eIF2α
Normalized fold change of p-mTOR
Normalized fold change of p-S6
37 °C
39 °C 4 h
39 °C 1 d
39 °C 4 d
p = 0.013
p = 0.023
p = 0.063
p = 0.002
p = 0.307
p = 0.606
p = 0.007
p = 0.495
p = 0.917
p = 0.003
p = 0.053
p = 0.341
p = 0.002
p = 0.307
p = 0.606
p = 0.358
p = 0.855
p = 0.238
p = 0.043
p = 0.051
p = 0.794
p = 0.025
p = 0.169
p = 0.947
p = 0.003
p = 0.174
p = 0.028

**Figure supplement 8. Differential effects of Hsp90 levels on eIF2α, mTOR, and S6.** Representative immunoblots of some translation-related proteins in A549 cells. β-actin serves as the loading control. The bar graphs show the quantitation of 3 biologically independent experiments, with densitometric scores determined from immunoblots. Values were normalized to the loading control β-actin. The phosphoproteins were normalized to the respective total proteins. The data are represented as mean values ± SEM. The statistical significance between the groups was analyzed by two-tailed unpaired Student's t-tests.

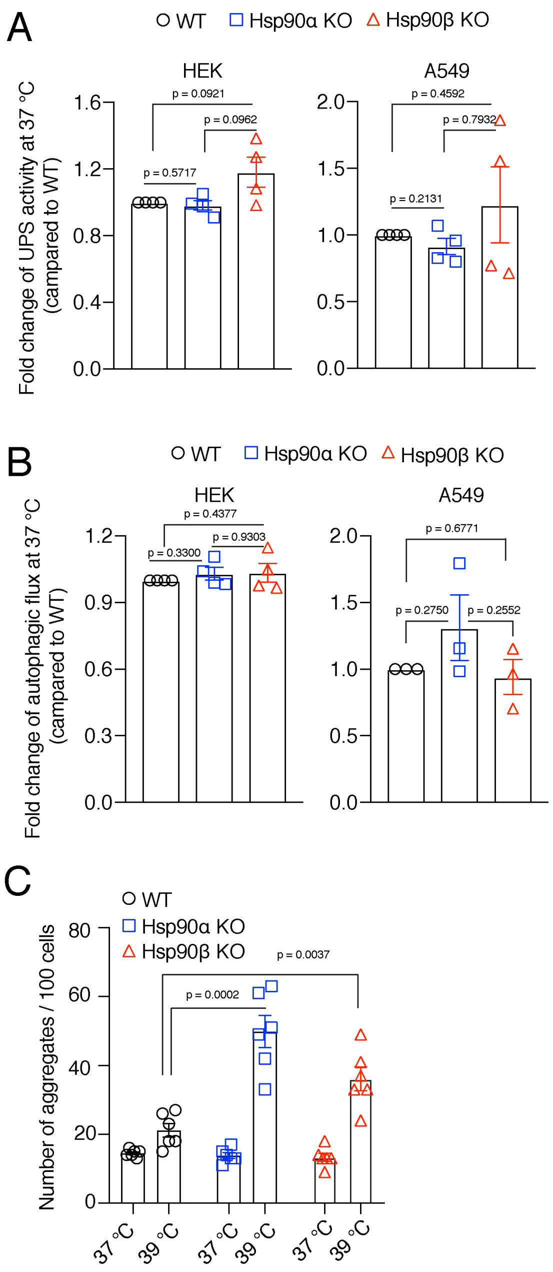


**Figure supplement 9. Hsp90α/β KO cells maintain WT levels of protein degradation activities in unstressed conditions, but have more protein aggregates. (A)** Flow cytometric determination of the *in vivo* UPS activity using the Ub-M-GFP and Ub-R-GFP reporter plasmids (n = 4 biologically independent samples). (**B**) Flow cytometric measurement of autophagic flux using a mCherry-GFP-LC3 reporter. Flux is calculated as the ratio of the mean fluorescence intensities of mCherry and GFP-positive cells (n = 4 biologically independent samples). **(C)** Quantitation of the number of EGFP-Q74 aggregates per 100 cells from representative micrographs such as the ones of Figure 7D. For each bar, data are from 6 different experiments, each normalized to 100 cells. The data are represented as mean values ± SEM for all bar graphs. The statistical significance between the groups was analyzed by two-tailed unpaired Student's t-tests.

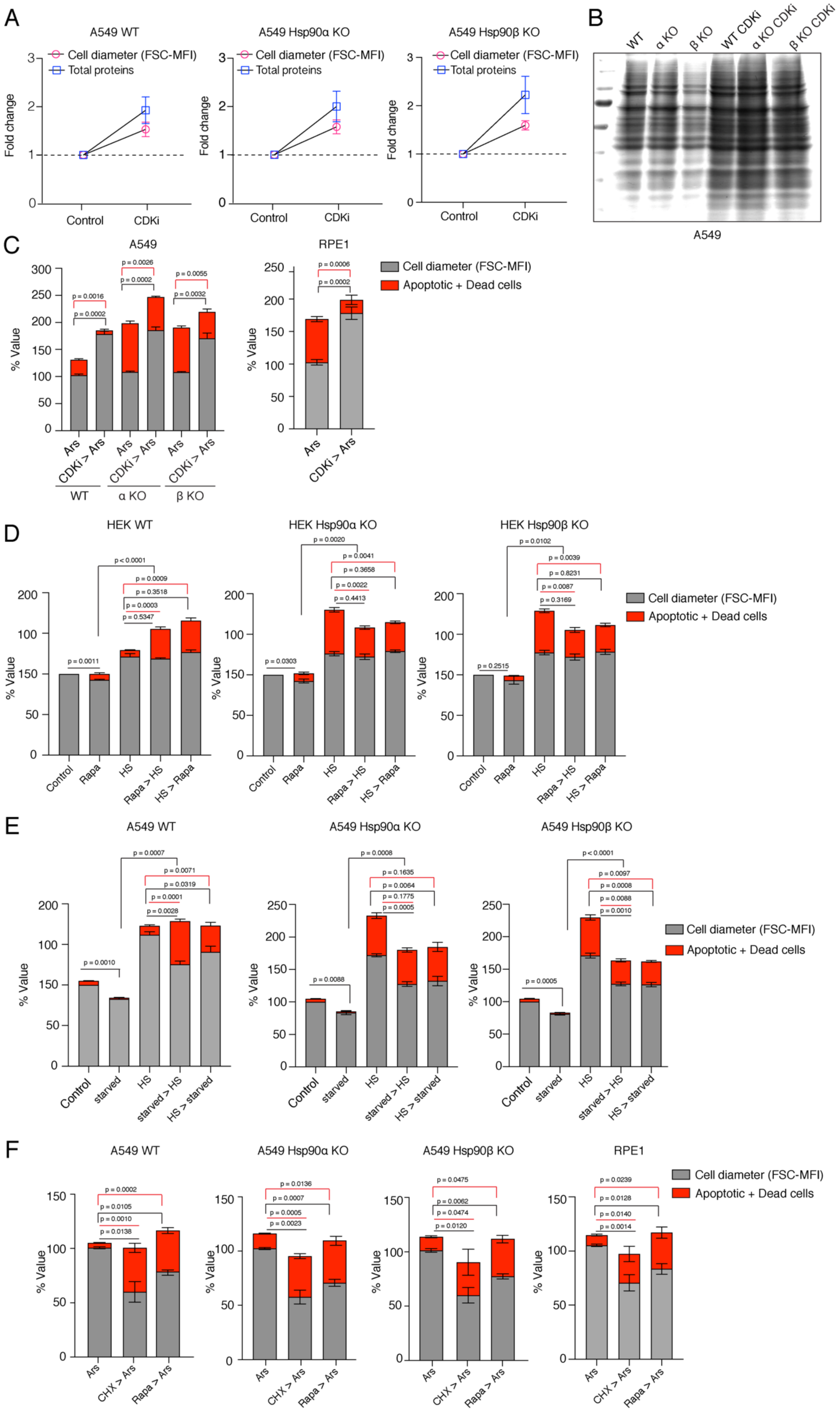
A
A549 WT
A549 Hsp90α KO
A549 Hsp90β KO
Cell diameter (FSC-MFI)
Total proteins
Fold change
Control
CDKi
B
WT
α KO
β KO
WT CDKi
α KO CDKi
β KO CDKi
A549
C
A549
RPE1
% Value
Ars
CDKi > Ars
WT
α KO
β KO
Cell diameter (FSC-MFI)
Apoptotic + Dead cells
D
HEK WT
HEK Hsp90α KO
HEK Hsp90β KO
% Value
Control
Rapa
HS
Rapa > HS
HS > Rapa
Cell diameter (FSC-MFI)
Apoptotic + Dead cells
E
A549 WT
A549 Hsp90α KO
A549 Hsp90β KO
% Value
Control
starved
HS
starved > HS
HS > starved
Cell diameter (FSC-MFI)
Apoptotic + Dead cells
F
A549 WT
A549 Hsp90α KO
A549 Hsp90β KO
RPE1
% Value
Ars
CHX > Ars
Rapa > Ars
Cell diameter (FSC-MFI)
Apoptotic + Dead cells

**Figure supplement 10. Smaller cells are more susceptible to additional stress.** **(A)** Cell size was enlarged by treating cells with 100 nM CDKi for 3 days. Fold change of cell size (represented by the FSC-MFI values) and total proteins (determined as MFI-FL1 values) were analyzed by flow cytometry. Cells were fixed, and total proteins were stained using Alexa Fluor 488 NHS ester (n = 3 biologically independent experiments). **(B)** Coomassie-stained protein gel showing protein samples from equal numbers of cells treated or not with CDKi (representative images of n = 2 biologically independent experiments). **(C)** Cell size was enlarged by treating cells with CDKi for 3 days, before cells were treated with sodium arsenite (40 μM) for 3 days. Cell size (% MFI) and cell death (% annexin V and PI-positive) are measured by flow cytometry. α KO, Hsp90α KO; β KO, Hsp90β KO cells. **(D)** Order of treatment experiment as in Figure 8C, but with HEK cells. Cell size (% MFI) and relative cell death (% annexin V and PI-positive) are quantified by flow cytometry. The values for cell size and death in the different experimental conditions are normalized to the respective 37 °C controls (n = 3 biologically independent experiments). **(E)** Cell size was reduced by serum starvation (starved) for 3 days before subjecting cells to chronic HS for 3 additional days (starved > HS). HS > starved, the two treatments were done the other way around (HS > starved). Cell size (% MFI) and relative cell death (% annexin V-PI positive) were quantified by flow cytometry and normalized to the respective 37 °C controls (n = 3 experimental samples). **(F)** Order of treatment experiment with CHX and rapamycin to reduce cell size first for 3 days and then subjecting cells to oxidative stress with 10 *μ*M sodium arsenite (Ars) for 1 day. Cell size (% MFI) and relative cell death (% annexin V and PI-positive) were quantified by flow cytometry and normalized to the Ars single treatment controls (n = 3 experimental samples).

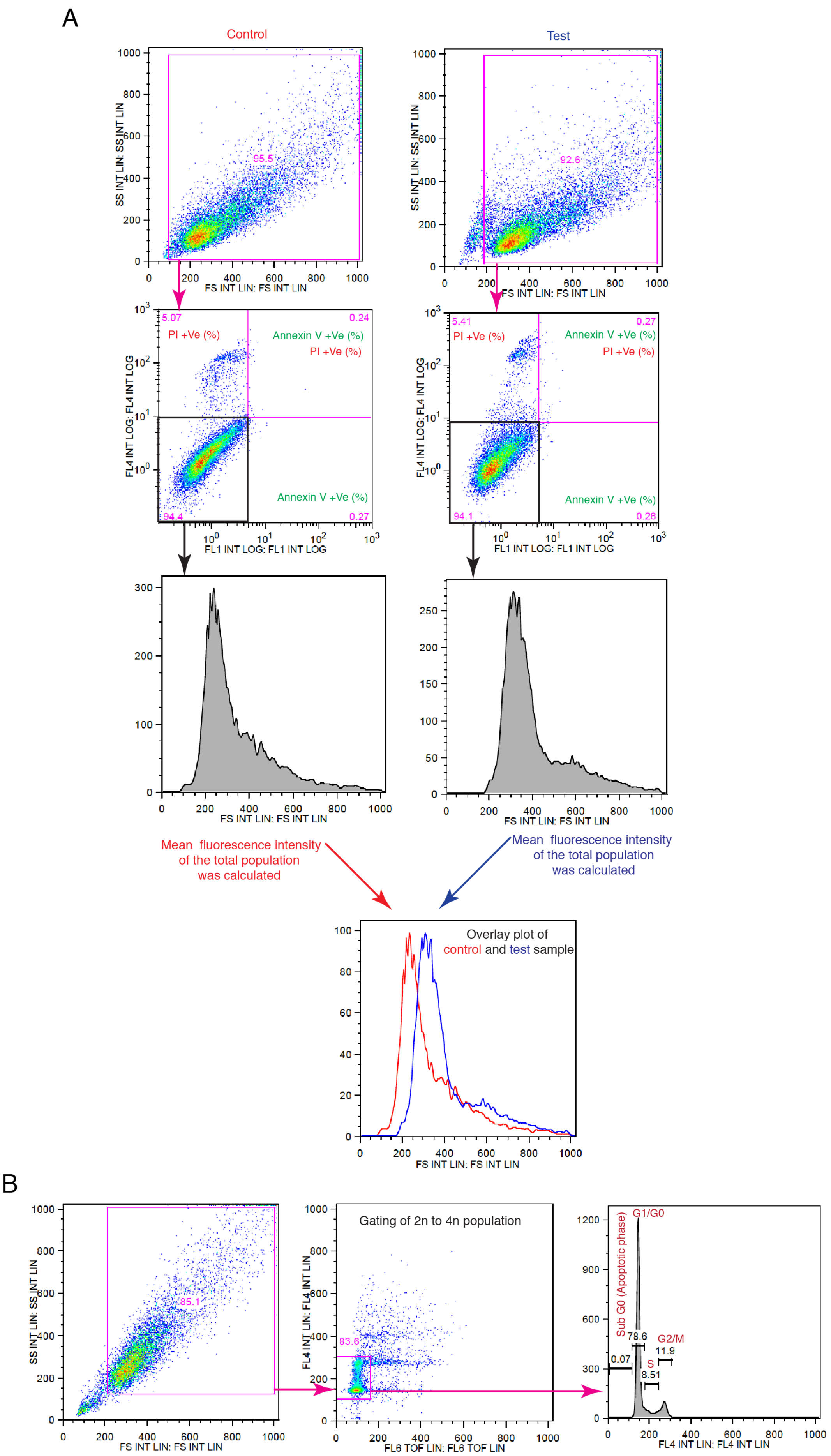
A
Control
Test
SS INT LIN: SS INT LIN
FS INT LIN: FS INT LIN
95.5
92.6
FL4 INT LOG: FL4 INT LOG
FL1 INT LOG: FL1 INT LOG
5.07
0.24
0.27
94.4
5.41
0.27
0.26
94.1
PI +Ve (%)
Annexin V +Ve (%)
PI +Ve (%)
Annexin V +Ve (%)
Mean fluorescence intensity of the total population was calculated
Mean fluorescence intensity of the total population was calculated
Overlay plot of control and test sample
B
85.1
Gating of 2n to 4n population
FL4 INT LIN: FL4 INT LIN
FL6 TOF LIN: FL6 TOF LIN
83.6
Sub G0 (Apoptotic phase)
G1/G0
G2/M
S
0.07
78.6
8.51
11.9

**Figure supplement 11. Schematic representation of the flow cytometric strategies for cell size and cell cycle analyses.** (**A**) Gating and strategy for cell size analyses relevant to Figures 1B, E, and H, 2C, 3E, 5B, 6E, 8B-C, and F, 10A and F, and Figure supplements 1C, 2B, 3D, 3G, 3J, 10A, and 10C-F. For size measurements, cell populations were gated based on the values of the forward scatter (FSC). (**B**) Gating and analysis strategy for cell cycle analyses relevant to Figures 1C-D, and I, 10B-C, and Figure supplement 1F.

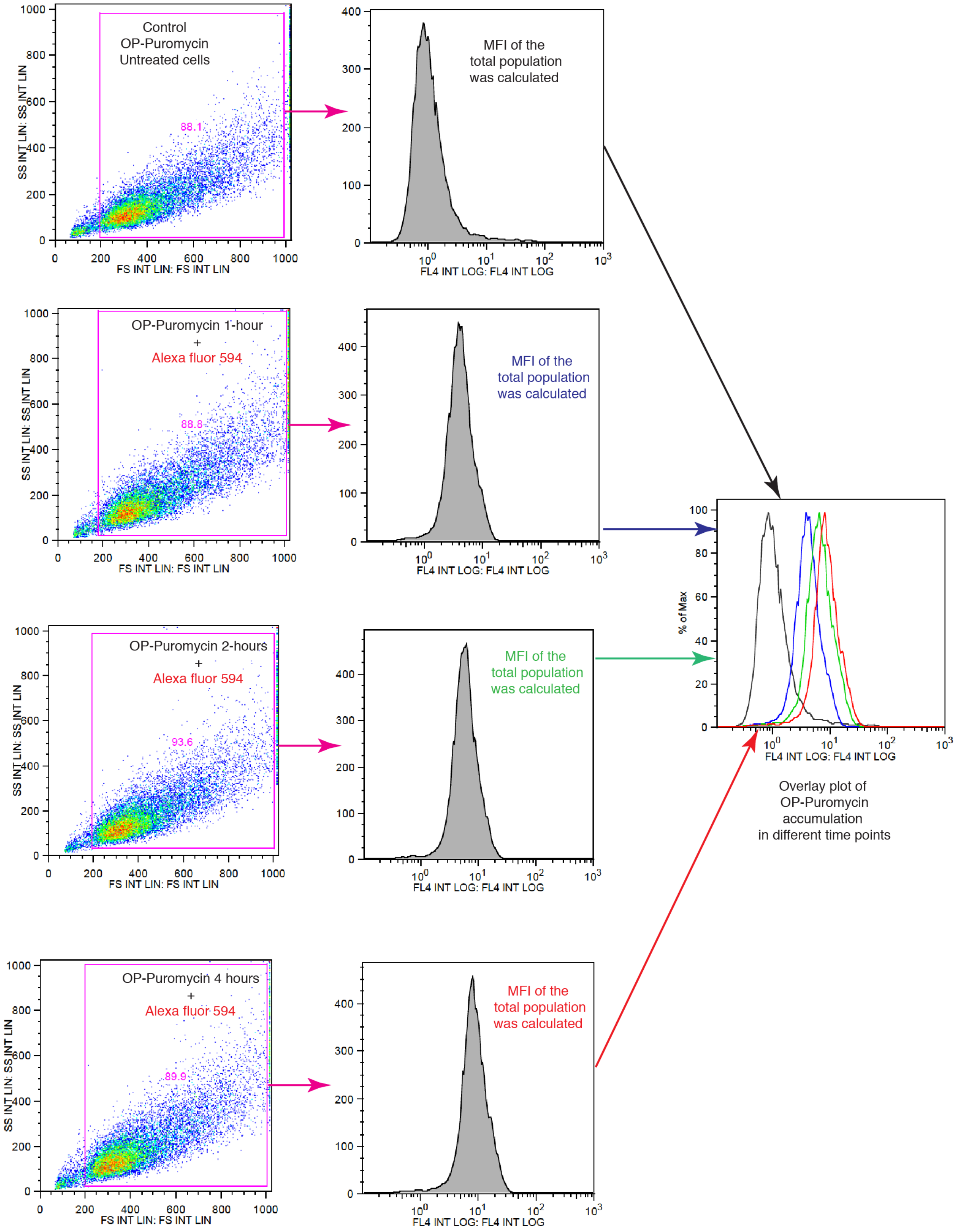


**Figure supplement 12. Schematic representation of the flow cytometric strategies to measure translation.** Relevant to Figure 6A and E, and Figure supplement 7A-B.

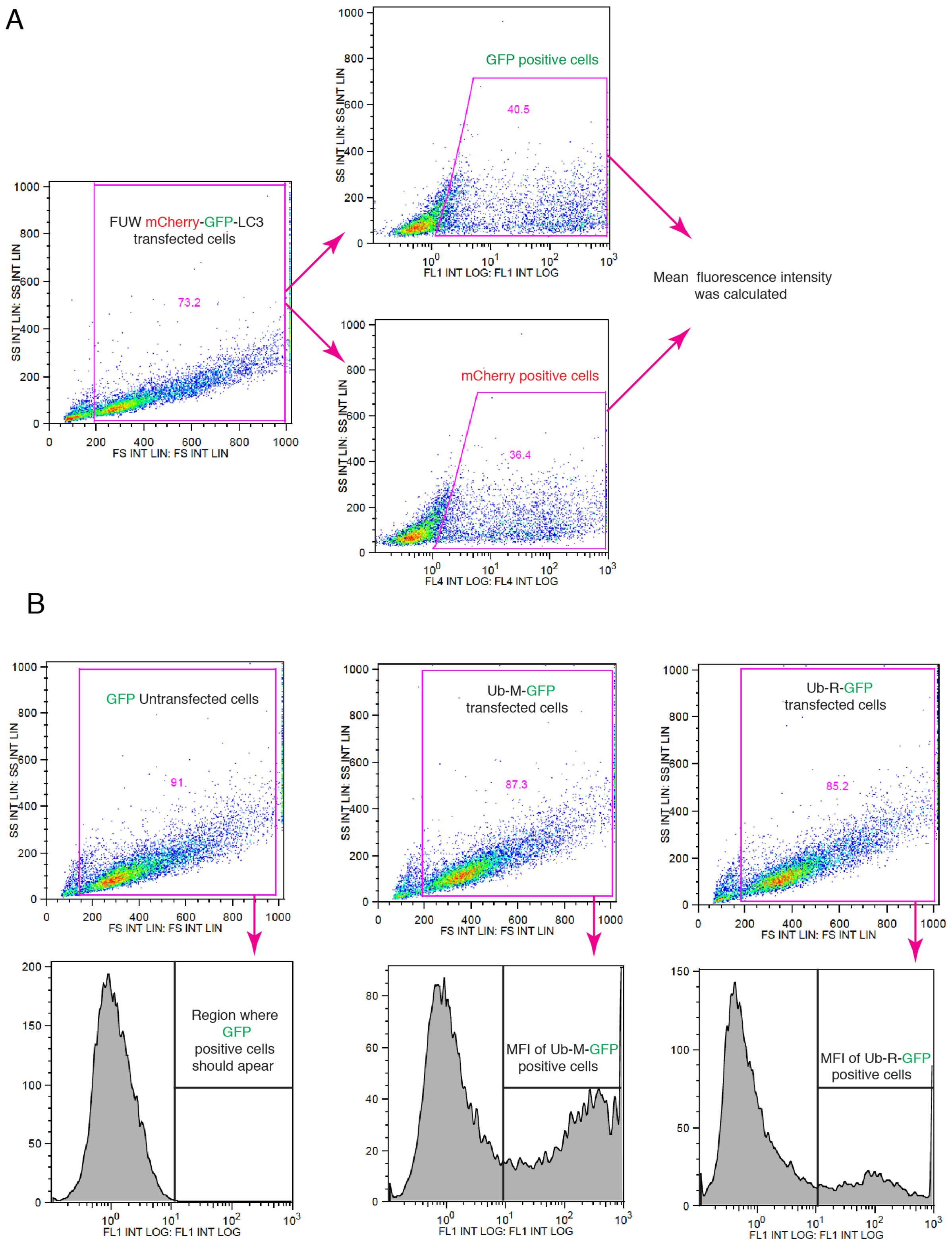


**Figure supplement 13. Schematic representation of the flow cytometric strategies for measuring autophagic flux and *in vivo* UPS activities**. (**A**) Gating strategy for autophagic flux measurements, relevant to Fig. 7B and Figure supplement 9B. (**B**) Gating strategy for measuring UPS activity; related to Figure 7A and Figure supplement 9A.

**Table 1. List of oligonucleotides used to generate the expression vectors for the shRNAs shHSF1-1 and shHSF1-2.**

| shRNA | Oligo name | Sequence 5' to 3' |
|---|---|---|
| shHSF1-1 | HSF1 shRNA 3UTR forward | 5'-CCG GGC AGG TTG TTC ATA GTC AGA ACT CGA GTT CTG ACT ATG AAC AAC CTG CTT TTT G-3' |
| | HSF1 shRNA 3UTR reverse | 5'-AAT TCA AAA AGC AGG TTG TTC ATA GTC AGA ACT CGA GTT CTG ACT ATG AAC AAC CTG C-3' |
| shHSF1-2 | HSF1 shRNA CDS forward | 5'-CCG GCC AGC AAC AGA AAG TCG TCA ACT CGA GTT GAC GAC TTT CTG TTG CTG GTT TTT G-3' |
| | HSF1 shRNA CDS reverse | 5'-AAT TCA AAA ACC AGC AAC AGA AAG TCG TCA ACT CGA GTT GAC GAC TTT CTG TTG CTG G-3' |

**Table 2. List of antibodies used in this study.**

| Antigen/Antibody | Supplier | Catalog # | Host | Clonality | Dilution |
|---|---|---|---|---|---|
| anti-GAPDH | HyTest Ltd. | 5G4 | Mouse | monoclonal | 1:1000 |
| anti-Hsp25/27 | StressMarq | SMC-114 | Mouse | monoclonal | 1:1000 |
| anti-Puromycin | Sigma-Aldrich | MABE343 | Mouse | monoclonal | 1:22000 |
| anti-Lamin B1 | Cell Signaling Technology | 12586 | Rabbit | monoclonal | 1:1000 |
| anti-Hsp40/Hdj1 | Enzo Lifesciences | ADI-SPA-400 | Rabbit | polyclonal | 1:1000 |
| anti-Hsf1 | Enzo Lifesciences | ADI-SPA-901 | Rabbit | polyclonal | 1:1000 |
| anti-Hsp70 | StressMarq | SMC-100 | Mouse | monoclonal | 1:1000 |
| anti-Phospho-eIF2α (Ser51) | Cell Signaling Technology | 3597 | Rabbit | monoclonal | 1:1000 |
| anti-eIF2α | Cell Signaling Technology | 9722 | Rabbit | polyclonal | 1:1000 |
| anti-Hsp90α (9D2) | Enzo Lifesciences | ADI-SPA-840 | Rat | monoclonal | 1:1000 |
| anti-Hsp90β (scFv H90-10) | Geneva Antibody Facility | ABCD_A0870 | Mouse | monoclonal | 1:2000 |
| anti-mTOR | Cell Signaling Technology | 2983 | Rabbit | monoclonal | 1:2000 |
| anti-Phospho-mTOR (Ser2448) | Cell Signaling Technology | 2971 | Rabbit | polyclonal | 1:1000 |
| anti-Phospho-S6 Ribosomal Protein (Ser235/236) | Cell Signaling Technology | 4858 | Rabbit | monoclonal | 1:1000 |
| anti-S6 Ribosomal Protein | GeneTex | GTX130450 | Rabbit | polyclonal | 1:1000 |
| anti-Hsc70 | StressMarq | SMC-151 | Mouse | monoclonal | 1:2000 |
| anti-beta-Actin | Cell Signaling Technology | 3700 | Mouse | monoclonal | 1:2000 |
| anti-4E-BP1(N1C3) | GeneTex | GTX109162 | Rabbit | polyclonal | 1:1000 |
| anti-phospho-4E-BP1(Thr37/46) | GeneTex | GTX133182 | Rabbit | polyclonal | 1:1000 |

**The following Supplementary Material is available as separate files:**

- **Source Data 1.** A subset of the proteomic data of WT and Hsp90α/β KO HEK293T cells. The main and first tab of the Excel file contains all of the unfiltered proteomic data. The following abbreviations (column headers) are used: 37C-WT, WT cells cultured at 37 °C; 39C-1d-WT, WT cells cultured at 39 °C for 1 day; 39C-4d-WT, WT cells cultured at 39 °C for 4 days; 37C-a-KO, Hsp90α KO cells cultured at 37 °C, 39C-1d-a-KO, Hsp90α KO cells cultured at 39 °C for 1 day; 39C-4d-a-KO, Hsp90α KO cells cultured at 39 °C for 4 days; 37C-b-KO, Hsp90β KO cells cultured at 37 °C; 39C-1d-b-KO, Hsp90β KO cells cultured at 39 °C for 1 day; 39C-4d-b-KO, Hsp90β KO cells cultured at 39 °C for 4 days. The other tabs contain the source data related to the figures indicated by the name of the tab; different colors are used to distinguish different the groups.
- **Source Data 2**. Original values related to all graphs. Different colors are randomly used to distinguish different the groups.
- **Source Data 3**. Original Western blots related to the main figures.
- **Source Data 4**. Original Western blots related to the figure supplements.
