## Supplementary figures and images for "Hsf1 and the molecular chaperone Hsp90 support a “rewiring stress response” leading to an adaptive cell size increase in chronic stress"

### Original Western blots related to the figure supplements.

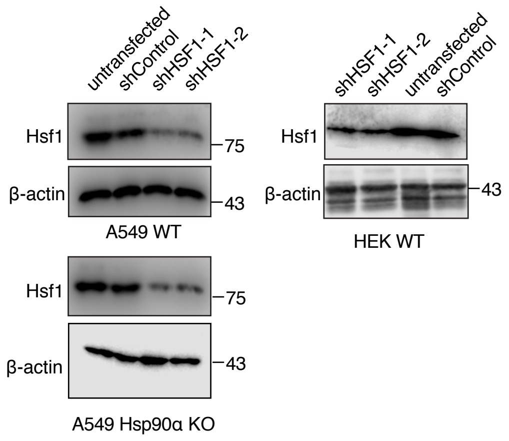

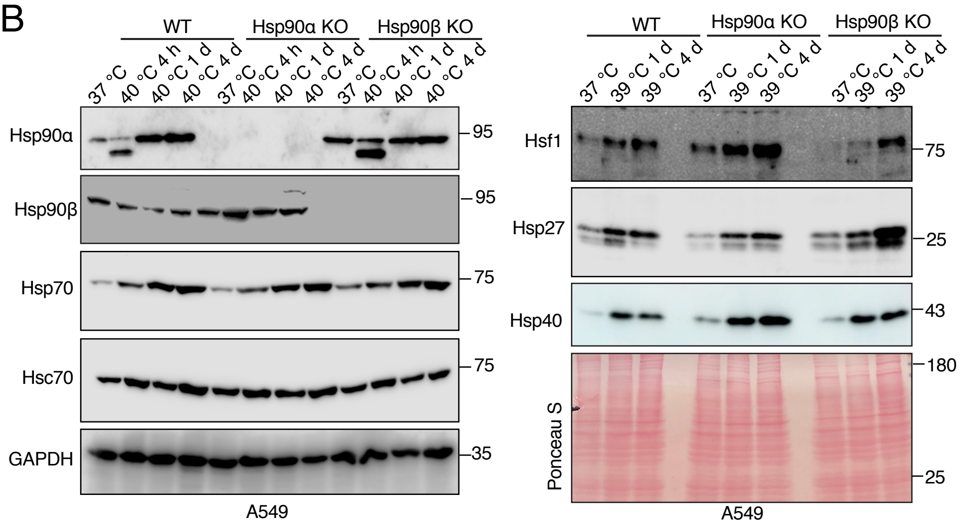

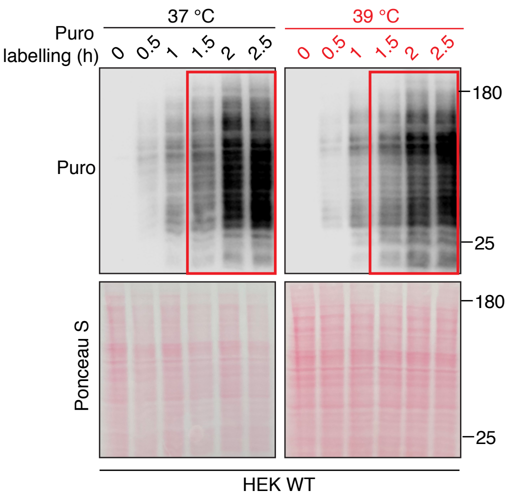

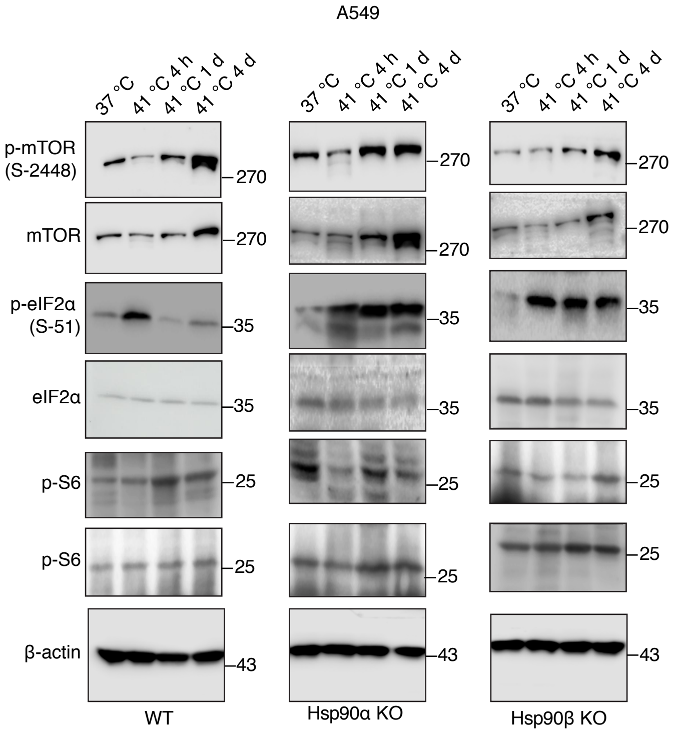

### Original Western blots related to the main figures

2A

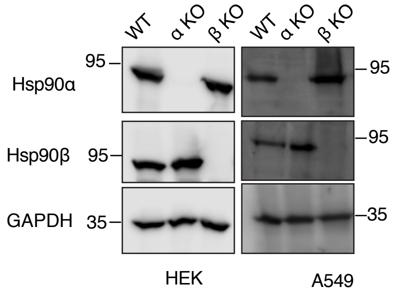


3B

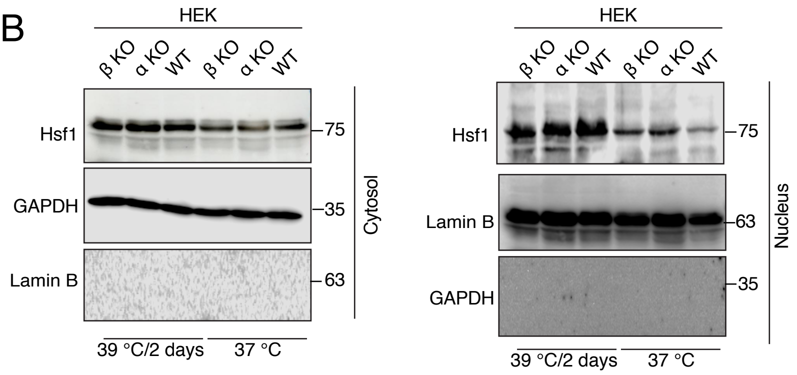


4B

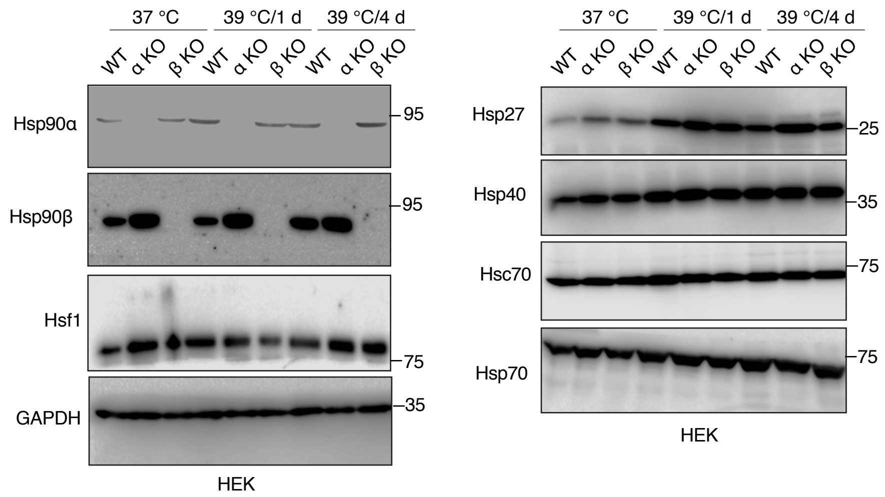


6C

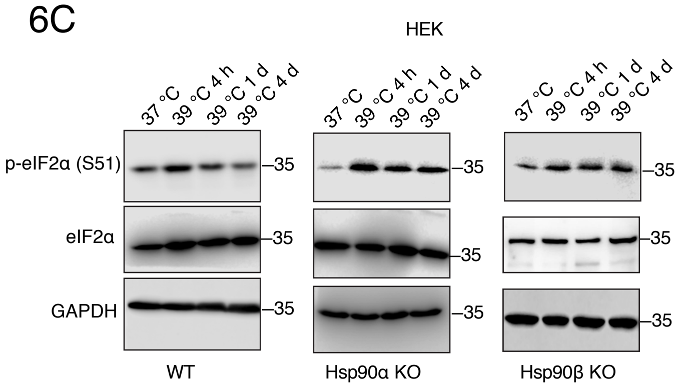


6D

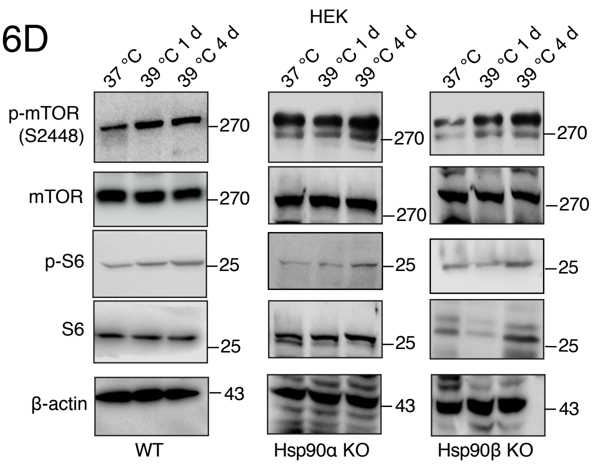


9

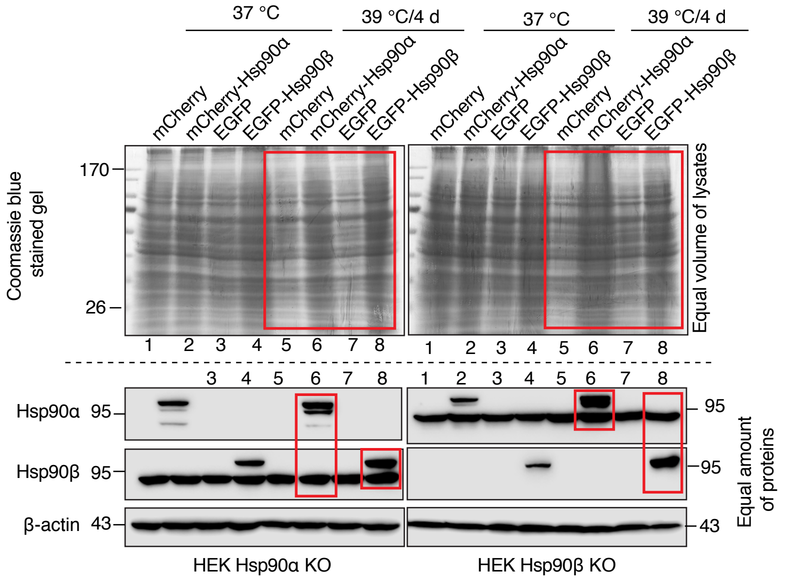
